# Perturbing neural representations of working memory with task-irrelevant interruption

**DOI:** 10.1101/716613

**Authors:** Nicole Hakim, Tobias Feldmann-Wüstefeld, Edward Awh, Edward K. Vogel

## Abstract

Working memory maintains information so that it can be used in complex cognitive tasks. A key challenge for this system is to maintain relevant information in the face of task-irrelevant perturbations. In this series of experiments, we investigated the impact of task-irrelevant interruptions on neural representations of working memory. We recorded electroencephalogram (EEG) activity in humans while they performed a working memory task. On a subset of trials, we interrupted participants with salient, but task-irrelevant objects. To track the impact of these task-irrelevant interruptions on neural representations of working memory, we measured two well-characterized, temporally sensitive EEG markers that reflect active, prioritized working memory representations: the contralateral delay activity (CDA) and lateralized alpha power (8-12hz). Following interruption, we found that CDA momentarily sustained, but was gone by the end of the trial. Lateralized alpha power was immediately influenced by the interrupters, but recovered by the end of the trial. This suggests that dissociable neural processes contribute to the maintenance of working memory information. Additionally, we found that task expectancy modulated the timing and magnitude of how these two neural signals responded to task-irrelevant interruptions, suggesting that the brain’s response to task-irrelevant interruption is shaped by task context. The distinct time courses of and influence of task context on these two neural signatures of working memory have many interesting theoretical implications about how information is actively maintained in working memory.

**Significance statement:** Working memory plays a central role in intelligent behaviors because it actively maintains relevant information that is easily accessible and manipulatable. In everyday life, we are often interrupted while performing such complex cognitive tasks. Therefore, understanding how working memory responds to and overcomes momentary task-irrelevant interruptions is critical for us to understand how complex cognition works. Here, we unveil how two distinct neural signatures of working memory respond to task-irrelevant interruptions by recording electroencephalogram activity in humans. Our findings raise long-standing theoretical questions about how different neural and cognitive processes contribute to the maintenance of information in working memory.

## Introduction

Working memory is a large-scale neural system that maintains readily accessible task-relevant information via active neural firing. A key challenge for this system is to protect these active representations from task-irrelevant interruptions. Extensive prior work has characterized how the presence of irrelevant information during the encoding of targets (distractors) impacts working memory representations (Postle et al., 2005; Clapp et al., 2010; Gaspar and McDonald, 2014; Feldmann-Wüstefeld and Vogel, 2018). This work has revealed that the presence of distractors during this initial encoding period (0-500ms) greatly reduces working memory performance, in part because these items compete with targets for limited representational space in working memory (Vogel et al., 2005). After this initial encoding period, presence of irrelevant information (interrupters) has a reduced, but still measurable impact on performance (Vogel et al., 2006). These interrupters have less of an impact because working memory representations have reached a more stable state, which is consistent with the time-course of neural measures of working memory representations (Vogel et al., 2006; Ikkai et al., 2010). This reduced impact is also likely due to the formation of concurrent visual long-term memory representations that represent the targets in a passive, yet still accessible format (e.g. Fukuda & Vogel, in press; Woodman and Chun, 2006; Chun and Turk-Browne, 2007). Together, these concurrent active and passive representations of targets reduce the behavioral impact of interruption during working memory maintenance. Yet, despite robust behavioral performance, current models of working memory still predict that onsets of task-irrelevant interruption should produce a momentary perturbation of the maintained target representations during which attention is withdrawn from the targets and at least temporarily applied to the positions of the interrupters (Bisley and Goldberg, 2003; Bisley et al., 2004). However, the consequences of such a brief withdrawal of attention on the neural signatures of working memory are not well understood. Here, we seek to measure the impact that task-irrelevant interruption has on the ongoing active neural representations of targets held in working memory.

To track the impact of task-irrelevant interruption on neural representations of working memory, we measured two well-characterized, temporally sensitive EEG markers that reflect active, prioritized working memory representations: the contralateral delay activity (CDA) and lateralized alpha power (8-12hz). The CDA is a sustained negative-going wave in human EEG that tracks current working memory load. It is sensitive to trial-by-trial fluctuations in working memory performance and distinguishes stable individual differences in working memory (Vogel and Machizawa, 2004; Luria et al., 2016). This component is thought to reflect an index of the current items that are actively represented in working memory (Feldmann-Wüstefeld et al., 2018; Hakim et al., 2018). Lateralized alpha power is similarly sensitive to task-relevant information. This signal is measured as a decrease in alpha power over posterior electrodes that are contralateral to the position of the attended items. However, despite its similarity to the CDA, it has been shown to be a distinct component of actively maintained information (Fukuda et al., 2015) that appears to primarily track the current position of spatial attention (Worden et al., 2000; Thut et al., 2006; Foster et al., 2016, 2017a). Topographic distributions of alpha power across the entire scalp have been shown to contain precise spatial information about remembered/attended stimuli (Foster et al., 2017a, 2017b), whereas lateralized alpha power have been used as an effective tool for establishing which visual hemifield is currently attended. Together, the CDA and lateralized alpha power respectively provide an item-based and space-based index of task-relevant information that is actively represented in working memory. Furthermore, because both signals are lateralized, we were able to isolate processing of the memory array by presenting the memory items laterally and the interrupters along the vertical midline of the display. As items on the vertical midline do not affect lateralized signals, the ongoing lateral measures only reflect processing of the memory representations. This allowed us to measure how these working memory representations respond to task-irrelevant interruption.

## Materials & Methods

### Overview of Experiments

In Experiment 1, we sought to determine how task-irrelevant interrupters impact ongoing working memory representations. We did this by presenting midline interrupters during the retention interval of a working memory task. In Experiment 2, we sought to determine whether the neural responses to task-irrelevant interrupters could be modulated by task expectancy. During all of these tasks, we recorded EEG activity from human participants.

## Experiment 1

### Participants

Twenty-two volunteers, naïve to the objective of the experiment participated for payment (∼15 USD per hour). The data of 2 participants were excluded from the analysis because of too many artifacts, poor behavioral performance (see below for criteria) or technical problems. The remaining 20 participants (12 male) were between the ages of 19-30 (M = 22.7, SD = 3.4). Participants in all experiments reported normal or corrected-to-normal visual acuity as well as normal color vision. All experiments were conducted with the written understanding and consent of each participant. The University of Chicago Institutional Review Board approved experimental procedures.

### Stimuli

All stimuli were presented on a gray background (∼33.3 cd/m^2^). Cue displays showed a small central fixation dot (0.2° × 0.2°). A horizontal diamond comprised of a green (RGB = 74, 183, 72; 52.8 cd/m^2^) and a pink (RGB = 183, 73, 177; 31.7 cd/m^2^) triangle appeared on the vertical midline 0.65° above the fixation dot. In 50% of the trials, the pink triangle pointed to the left side and the green triangle pointed to the right side, in the remaining 50% of the trials this was inverse. Half the participants were instructed to attend the hemifield that the pink triangle pointed to, and the other half was instructed to attend the hemifield the green triangle pointed to. Memory displays showed a series of colored squares (1.1° by 1.1°, mean luminance 43.1 cd/m^2^). Colors for the squares were selected randomly from a set of 11 possible colors (Red = 255 0 0; Green = 0 255 0; Blue = 0 0 255; Yellow = 255 255 0; Magenta = 255 0 255; Cyan = 0 255 255; Purple = 102 0 102; Brown = 102 51 0; Orange = 255 128 0; White = 255 255 255; Black = 0 0 0). Squares could appear within an area of the display subtending 3.5° to the left or right of fixation and 3.1° above and below fixation. There was the same number of squares in each hemisphere. Within each hemisphere, squares were as equally distributed between the upper and lower hemifields as possible. The interruption display showed four colored squares of the same size as the memory display, drawn from the remaining colors. These interrupting items were shown on the vertical midline with a randomly jittered horizontal offset of maximally 0.55° (half of an object). Retention interval displays were blank with a small central fixation dot (0.2° × 0.2°). Probe displays showed one colored square in each hemisphere in the same location as one of the squares in the original array. In 50% of the trials, the color was identical (no change trials) to the memory display. In the remaining 50% of the trials, it was one of the colors not used in the memory or interruption display (change trials). The same stimuli were used in all experiments.

### Apparatus

Participants were seated with a chin-rest in a comfortable chair in a dimly lit, electrically shielded and sound attenuated chamber. Participants responded with button presses on a standard keyboard that was placed in front of them. Stimuli were presented on an LCD computer screen (BenQ XL2430T; 120 Hz refresh rate; 61 cm screen size in diameter; 1920 × 1080 pixels) placed at 74 cm distance from participants. An IBM-compatible computer (Dell Optiplex 9020) controlled stimulus presentation and response collection.

### Procedure

Each trial began with a cue display (500 ms) indicating the relevant side of the screen (left or right). A memory display consisting of 6 colored squares in each hemisphere followed the cue display for 150 ms. Participants were instructed to memorize as many colored squares in the memory display from the cued side and to ignore the other side entirely, as that side would never be probed. Participants had to remember the items for a retention interval of 1,650 ms during which a central fixation dot was shown. In 25% of the trials, an interruption display appeared 500 ms after memory display offset for 150 ms. The total length of the retention interval was 1,650 ms, regardless of whether an interruption appeared. Participants were instructed to always ignore interruption displays. After the retention interval, a probe display appeared until response. Participants had to indicate whether the color at the probed location changed color (“?/” key) or did not change color (“z” key). After participants responded, the trial concluded and the next trial started after a blank inter-trial interval of 750 ms. Participants completed a total of 1200 trials (15 blocks of 80 trials), i.e. 300 trials with interruption and 900 trials without interruption. Information about average performance and a minimum break of 30 seconds was provided after each block. See Figure 1 for a visual depiction of the task.

**Figure 1.**
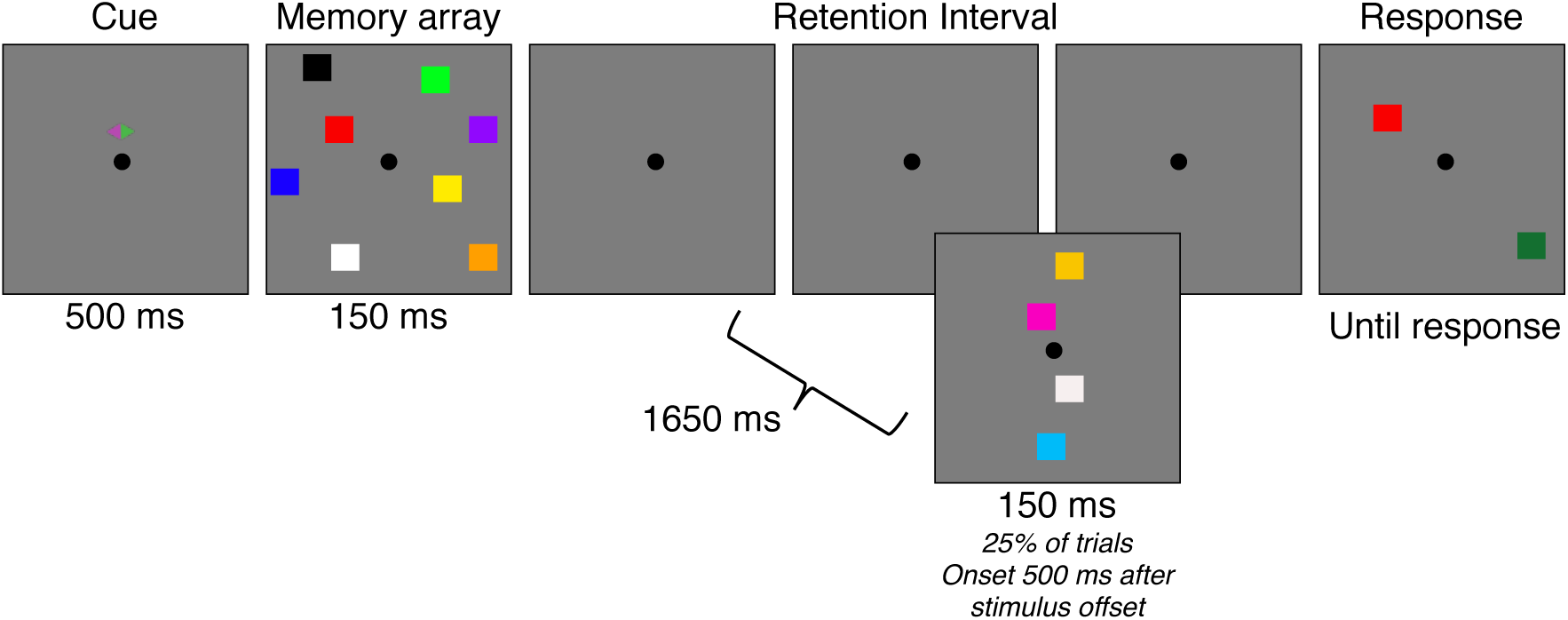
Task for Experiment 1. At the start of each trial, a cue appeared on the screen for 500 ms, which cued participants to attend one side of the screen. Then, an array of 4 colored squares briefly appeared (150 ms). On 25% of trials, during the blank retention interval (1650 ms), four colored squares appeared on the midline for 150 ms. Participants were told to always ignore the squares that appeared during the retention interval. After the retention interval, a response screen appeared with one square in each hemifield. Participants were told to report whether the square on the attended side was the same color as the original square in that location.

We presented the interrupters in locations that did not overlap with the locations of the memory items to avoid visual masking. Importantly, the relative position of interrupters and targets matter in lateralized change detection tasks. When interrupters are presented laterally with targets on the vertical midline, the neural signature of sustained interrupters suppression can be isolated (CDAp). Conversely, when interrupters are presented on the vertical midline and targets are presented laterally, the neural signature of target processing can be isolated (Feldmann-Wüstefeld & Vogel, 2018). Accordingly, since we were interested in how neural representations of targets are affected by interruption, we placed the interrupters along the vertical midline. Thus, reductions in CDA amplitude can be interpreted as dropping memory items, and reductions in lateralized alpha power can be interpreted as a shift of attention away from the laterally presented memory arrays.

### Behavioral Data Analysis

We separately analyzed performance for the trials with and without interruption. Performance was converted to a capacity score, K, calculated as N × (H-FA), where N is the set-size, H is the hit rate, and FA is the false alarm rate (Cowan, 2011). To compare performance between the two conditions, we used a two-tailed, repeated measures t-test

### Artifact Rejection

We recorded EEG activity from 30 active Ag/AgCl electrodes (Brain Products actiCHamp, Munich, Germany) mounted in an elastic cap positioned according to the International 10-20 system [Fp1, Fp2, F7, F8, F3, F4, Fz, FC5, FC6, FC1, FC2, C3, C4, Cz, CP5, CP6, CP1, CP2, P7, P8, P3, P4, Pz, PO7, PO8, PO3, PO4, O1, O2, Oz]. FPz served as the ground electrode and all electrodes were referenced online to TP10 and re-referenced off-line to the average of all electrodes. Incoming data were filtered [low cut-off = .01 Hz, high cut-off = 80 Hz, slope from low- to high-cutoff = 12 dB/octave] and recorded with a 500 Hz sampling rate. Impedances were kept below 10kΩ. To identify trials that were contaminated with eye movements and blinks, we used electrooculogram (EOG) activity and eye tracking. We collected EOG data with 5 passive Ag/AgCl electrodes (2 vertical EOG electrodes placed above and below the right eye, 2 horizontal EOG electrodes placed ∼1 cm from the outer canthi, and 1 ground electrode placed on the left cheek). We collected eye-tracking data using a desk-mounted EyeLink 1000 Plus eye-tracking camera (SR Research Ltd., Ontario, Canada) sampling at 1,000 Hz. Usable eye-tracking data were acquired for 20 out of 22 participants in Experiment 1 and 29 out of 30 participants in Experiment 2.

EEG was segmented off-line with segments time-locked to memory display onset. Eye movements, blinks, blocking, drift, and muscle artifacts were first detected by applying automatic detection criteria to each segment. After automatic detection (see below), trials were manually inspected to confirm that detection thresholds were working as expected. Participants were excluded if they had less than 100 correct trials remaining in any of the conditions. For the participants used in analyses, we rejected on average 21% of trials in Experiment 1, and 39% of trials in Experiment 2.

### Eye movements

We used a sliding window step-function to check for eye movements in the HEOG and the eye-tracking gaze coordinates. For HEOG rejection, we used a split-half sliding window approach. We slid a 100 ms time window in steps of 10 ms from the beginning to the end of the trial. If the change in voltage from the first half to the second half of the window was greater than 20 µV, it was marked as an eye movement and rejected. For eye-tracking rejection, we applied a sliding window analysis to the x-gaze coordinates and y-gaze coordinates (window size = 100 ms, step size = 10 ms, threshold = 0.5° of visual angle).

### Blinks

We used a sliding window step function to check for blinks in the VEOG (window size = 80 ms, step size = 10 ms, threshold = 30 µV). We checked the eye-tracking data for trial segments with missing data-points (no position data is recorded when the eye is closed).

### Drift, muscle artifacts, and blocking

We checked for drift (e.g. skin potentials) by comparing the absolute change in voltage from the first quarter of the trial to the last quarter of the trial. If the change in voltage exceeded 100 µV, the trial was rejected for drift. In addition to slow drift, we checked for sudden step-like changes in voltage with a sliding window (window size = 100 ms, step size = 10 ms, threshold = 100 □V). We excluded trials for muscle artifacts if any electrode had peak-to-peak amplitude greater than 200 µV within a 15 ms time window. We excluded trials for blocking if any electrode had at least 30 time-points in any given 200-ms time window that were within 1V of each other.

### Contralateral Delay Activity Analysis

Segmented EEG data was baselined from 200 ms to 0 ms before the onset of the memory displays. Artifact-free EEG segments from correct trials were averaged separately for each condition (no interruption, interruption) and separately for electrodes ipsi- and electrodes contralateral to the attended side. Then the difference between contra- and ipsi-lateral activity for the electrode pair PO7/PO8 was calculated (i.e., the CDA), resulting in two average waveforms for each participant. The average CDA amplitude was calculated for three time windows: before interruption onset (450-650 ms), after interruption onset (800-1,000 ms), before probe onset (1,300-1,500 ms). We then compared the CDA for each time window with a repeated measures two-tailed t-test. To measure the robustness of the CDA for each condition (reliable difference between contra- and ipsilateral activity), we also ran a one-sample t-test (against zero) for each time window.

### Lateralized Alpha Power Analysis

The same EEG segments as the CDA analysis were used in this analysis, however, the segments were not baselined. The raw EEG signal was band-pass filtered in the alpha band (8-12 Hz) using a two-way least-squares finite-impulse-response filter (“eegfilt.m” from EEGLAB Toolbox). Instantaneous power was then extracted by applying a Hilbert transform (‘hilbert.m’) to the filtered data. The resulting data were averaged separately for each condition (no interruption, interruption) and each laterality (contra-versus ipsi-lateral to cued hemifield) for the electrode pair PO7/PO8, resulting in four average waveforms for each participant. The average alpha power was calculated for the same three time windows as the CDA analysis. We then compared lateralized alpha power suppression for each time window with a repeated-measures two-tailed t-test. To measure the robustness of latearlized alpha power suppression for each condition (reliable difference between contra- and ipsilateral activity), we also ran a one-sample t-test (against zero) for each time window.

## Results

### Behavior

Performance (Figure 2), as measured by K, was significantly worse on trials that were interrupted (M=1.6), than on trials that were not interrupted (M=1.2), significant two-way repeated measures t-test, t(19)=4.428, p<0.001, M=0.408, 95% CI (0.215, 0.601).

**Figure 2.**
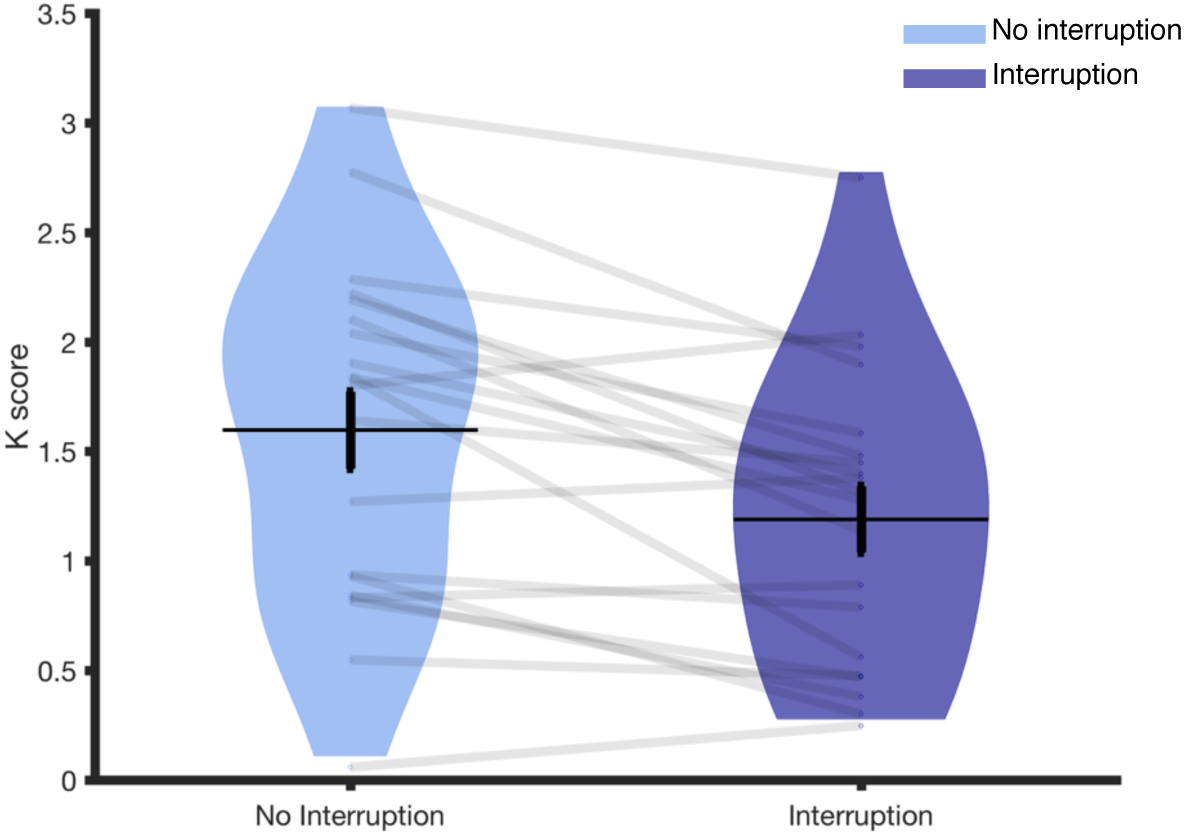
Behavioral performance for Experiment 1. Average performance is represented with the horizontal black line. Standard error is the vertical black line. The distribution of K scores for all participants is represented by the violin plot. Dots and light gray lines represent one participant’s performance.

### Contralateral Delay Activity

#### Pre-Interruption (450-60 ms)

The CDA (Figure 3) was robust before interruption onset (450-650 ms) on trials with, t(19)=-3.187, p=0.005, M=-0.707, 95% CI (−1.171 - 0.243), and without, t(19)=-4.053, p=0.001, M=-0.837, 95% CI (−1.270 -0.405) interruption. There was not a significant difference in CDA amplitude between trials with and without interruption during this time window, t(19)=-1.394, p=0.179, M=-0.131, 95% CI (−0.327, 0.066).

**Figure 3.**
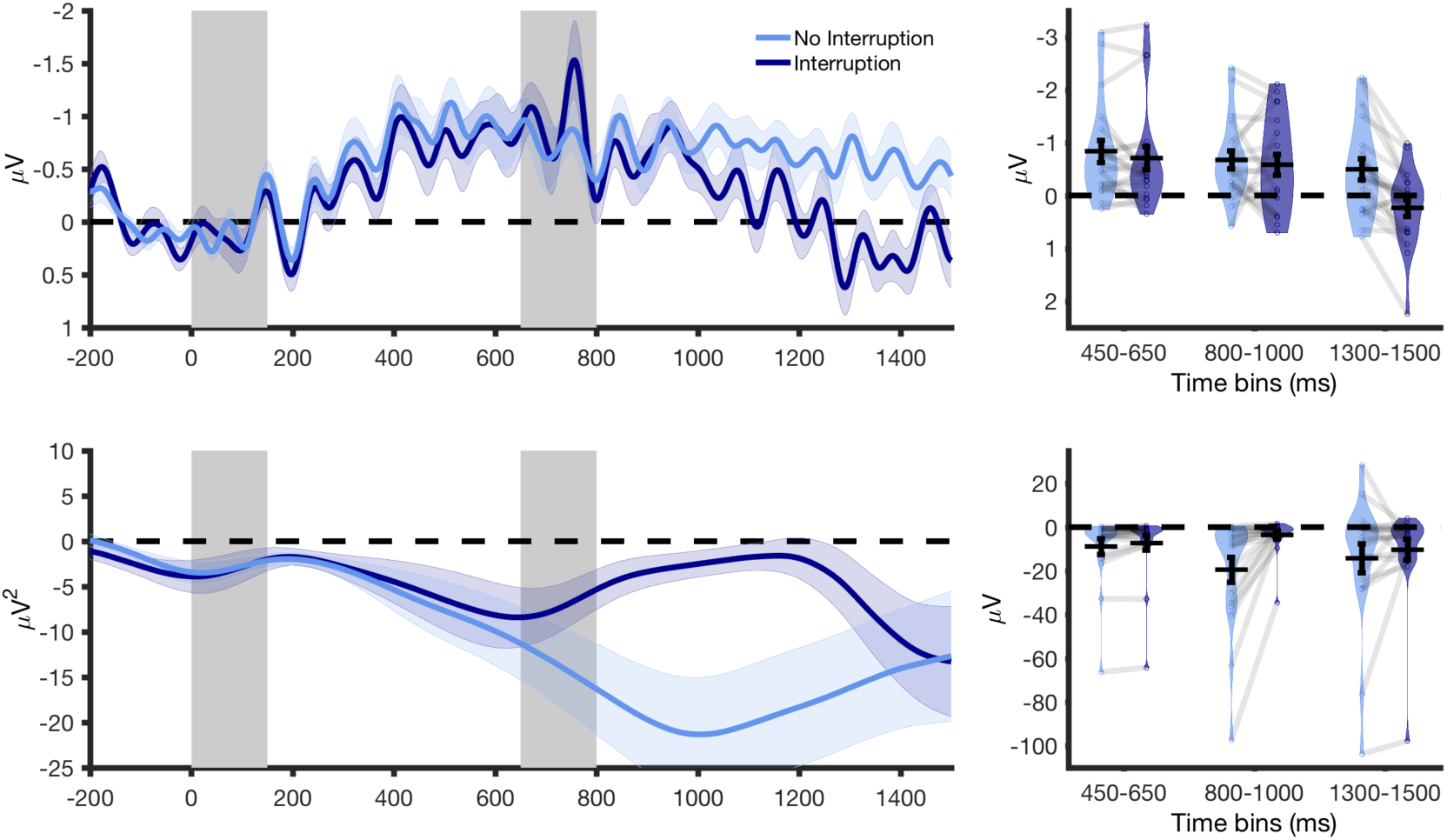
Contralateral delay activity and lateralized alpha power for Experiment 1. (Top) CDA amplitude over time. The first gray bar (timepoint 0-150) in both graphs represents when the memory array was on the screen. The second gray bar (timepoint 650-800) represents when the interrupters were on the screen in the Interruption condition. Light colored envelopes around each line represent standard error. (Bottom) Lateralized alpha power over time.

#### Post-Interruption I (800-1,000 ms)

Immediately following the offset of the interruption (800-1,000 ms), the CDA remained robust for both conditions, No Interruption: t(19)=-4.016, p=0.001, M= -0.674, 95% CI (−1.025 -0.323); Interruption: t(19)=-2.928, p=0.009, M=-0.583, 95% CI (−1.000 -0.166). Again, there was not a significant difference in CDA amplitude between trials with and without interruption, t(19)= -0.525, p= 0.606, M=-0.091, 95% CI (−0.452 0.271).

#### Post-interruption II (1,300-1,500 ms)

By the end of the trial (1,300-1,500 ms), however, there was a significant difference in CDA amplitude between trials with and without interruption, t(19)= -5.145, p= <0.001, M=-0.731, 95% CI (-1.028, -0.434). On trials without interruption, the CDA remained robust, t(19)=-2.535, p=0.020, M= -0.496, 95% CI (-0.906 -0.086). However, on trials with interruption, the CDA was no longer significantly different from zero, 1.445, p=0.165, M=0.234, 95% CI (-0.105 0.574).

### Lateralized Alpha Power

#### Pre-Interruption (450-60 ms)

Alpha power (Figure 3) was significantly more negative at contralateral compared to ipsilateral electrodes before interruption onset (450-650 ms) on trials with, t(19)=-2.131, p=0.046, M=-7.264, 95% CI (−14.398 -0.130), and without interruption, t(19)=-2.517, p=0.021, M=-8.815, 95% CI (−16.145 -1.486). Alpha power was significantly more lateralized on trials without interruption than trials with interruption during this time window, t(19)= -2.573, p= 0.019, M= -1.551, 95% CI (−2.813 -0.289). We suspect that this may be due to time smearing in the alpha analysis. Time smearing is a side-effect of Fourier transforms, as the calculation of power at any time point incorporates data from time points before and after the time point of interest. Therefore, the effect of the interruption may be smeared across time, causing it to appear like there are differences in alpha power before interruption onset when there actually are only differences after interruption onset.

#### Post-Interruption I (800-1,000 ms)

Immediately following the offset of the interruption (800-1,000 ms), lateralization of alpha power remained robust following trials without interruption, t(19)=-3.423, p=0.003, M= -19.393, 95% CI (−31.250 -7.536). However, alpha power was not significantly lateralized following trials with interruption, but this effect was trending towards significance, t(19)=-2.054, p=0.054, M=-3.554, 95% CI (−7.175 0.067). During this time window, lateralized alpha power was significantly more lateralized on trials without interruption than trials with interruption, t(19)= -3.629, p= 0.002, M=-15.839, 95% CI (−24.974 -6.704

#### Post-interruption II (1,300-1,500 ms)

By the end of the trial (1,300-1,500 ms), however, there was no longer a significant difference in lateralized alpha power between the two conditions, t(19)= -0.904, p= 0.377, M= -3.879, 95% CI (−12.863 5.104). Lateralized alpha power was significantly different lateralized in both conditions, No Interruption: t(19)=-2.144, p=0.045, M= -14.207, 95% CI (−28.077 -0.337), Interruption: t(19)=-2.137, p=0.046, M=-10.328, 95% CI (−20.444 -0.213).

### Conclusions

In Experiment 1, participants’ working memory performance was reduced when they were interrupted during the retention interval. In addition, both the CDA and lateralized alpha power were negatively impacted by the interrupters, but this effect had distinct time courses for the two signals. The CDA briefly sustained following interruption while alpha power immediately became less lateralized. By the end of the trial, CDA was no longer present, but alpha power re-lateralized. These results suggest that the CDA and lateralized alpha power respond distinctly to task-irrelevant interruptions.

## Experiment 2

In Experiment 2, we sought to determine whether the neural responses to task-irrelevant interruptions could be modulated by task expectancy, or if they are fixed responses to interruption irrespective of the subject’s expectations. Thus, in Experiment 2, we compared the same 25% interruption condition employed in Experiment 1 with one in which interrupters were presented on 75% of trials. We predicted that a higher frequency of task-irrelevant interruptions should allow participants to be better prepared for interruptions. Accordingly, CDA and lateralized alpha power should sustain for longer following interruption. Importantly, we will also examine the onset and offset of the CDA and lateralized alpha power to examine whether the time course of the two subprocesses may be affected differently.

## Materials & Methods

### Participants

Thirty novel volunteers, naïve to the objective of the experiment participated for payment (∼15 USD per hour). The data of 10 participants were excluded from the analysis because of too many artifacts, poor behavioral performance or technical problems (same criteria as in Experiment 1). The remaining 20 participants (11 male) were between the ages of 19-32 (M = 23.54, SD = 3.85).

### Stimuli & Procedures

Stimuli were identical to Experiment 1 (Figure 1). Procedure was also identical to Experiment 1, except for the following changes. The retention interval was increased to 2000 ms. The experiment was divided in two halves in each of which the probability of interruption was varied. The order of the halves was counterbalanced across participants. The probability for interruption was 25% in one part (no interruption: 75%) and 75% in the other part (no interruption: 25%). This resulted in 2 × 2 design with the factors Interruption (no interruption versus interruption) and Probability (high versus low). Participants completed 1,920 trials in total (24 blocks of 80 trials each), 240 trials for each of the two low probability conditions and 720 trials for each of the two high probability conditions.

### Analysis

Behavioral and EEG data were analyzed analogously to Experiment 1, but included the additional factor Probability. For statistical analyses, we forwarded the mean CDA amplitude (contra-minus ipsilateral activity) to a two-way ANOVA with the within-subjects factors Interruption (interruption versus no interruption) and Probability (75% probability for interruption versus 25%). Additionally, for the CDA and lateralized alpha analyses, the time window before probe onset was 1800-2000 ms, as we extended the retention interval by 500 ms.

## Results

### Behavior

Performance (Figure 4), as measured by K, was significantly worse on trials that were interrupted (Low Probability M = 1.4, High Probability M = 1.6) than on trials that were not interrupted (Low Probability M = 1.6, High Probability M = 1.7), regardless of probability, significant main effect of Interruption, F(1,19)=21.288, p<0.001, η_p_^2^ =0.528.

**Figure 4.**
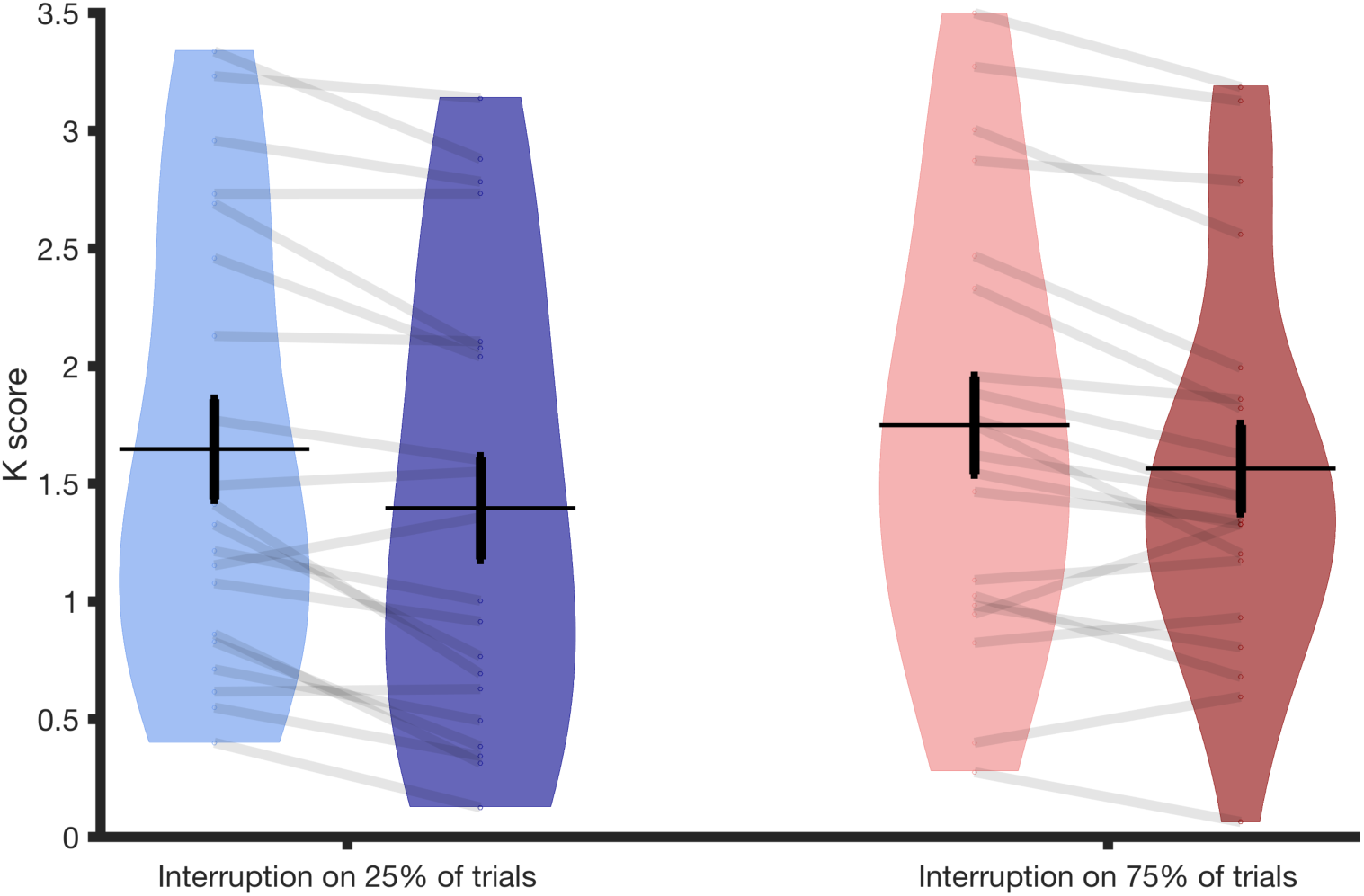
Behavioral performance for Experiment 2. Average performance is represented with the horizontal black line. Standard error is the vertical black line. The distribution of K scores for all participants is represented by the violin plot. Dots and light gray lines represent one participant’s performance.

### Contralateral Delay Activity

#### Pre-interruption II (450-650 ms)

Before interruption onset (450-650 ms), there was a significant CDA in all four conditions, all one-sample t-tests p≤0.002. Additionally, there was no difference in CDA amplitude between any of the conditions, p≥0.060 for the main effects of Interruption, Probability and their interaction. Though the interaction of Interruption and Probability was trending towards significance, p=0.060, η_p_^2^ =0.173.

#### Post-interruption I (800-1,000 ms)

Immediately following interruption offset (800-1,000 ms), the influence of interruption on CDA amplitude depended on the probability of being interrupted, significant interaction of Probability and Interruption: F(1,19)=9.951, p=0.005, η_p_^2^ = 0.344. Follow-up t-tests run separately for trials with and without interruption revealed that when interrupters were present, the amplitude of the CDA depended on the probability of interruption, t(19)=2.252, p=0.036. The CDA was significantly larger in the High Probability condition (M = -0.660) than in the Low Probability condition (M = -0.265). On trials without interruption, there was no difference in CDA amplitude between probabilities, t(19)=-0.858, p=0.402. The main effects of Interruption and Probability were not significant, p≥0.129.

#### Post-interruption II (1,800-2,000 ms)

By the end of the trial (1,800-2,000 ms), there was no difference in CDA amplitude between any of the conditions, p≥0.142 for the main effects of Interruption, Probability, and their interaction. There was no longer a significant CDA in any condition, all one-way t-tests p ≥ 0.088. CDA tends to decline over time, and by extending the delay period compared to Experiment 1, we may have reached the point at which the CDA tends to decline naturally (Vogel and Machizawa, 2004).

### Lateralized Alpha Power

#### Pre-interruption II (450-650 ms)

Alpha power (Figure 5) was significantly suppressed in all conditions before the interruption onset (450-650 ms), all one-sample t-tests p ≤ 0.005. The influence of interruption on alpha power suppression depended on the probability of interruption, significant interaction of Probability and Interruption: F(1,19)=4.881, p=0.040, η_p_^2^ = 0.204. As in Experiment 1, this pre-interruption difference could be due to time smearing of alpha power. There was no difference in lateralized alpha power between trials that were and were not interrupted for either the High (f(19)=-2.046, p=0.055) or Low (f(19)=0.279, p=0.789) probability trials, though this effect was trending in the High probability trials.

**Figure 5.**
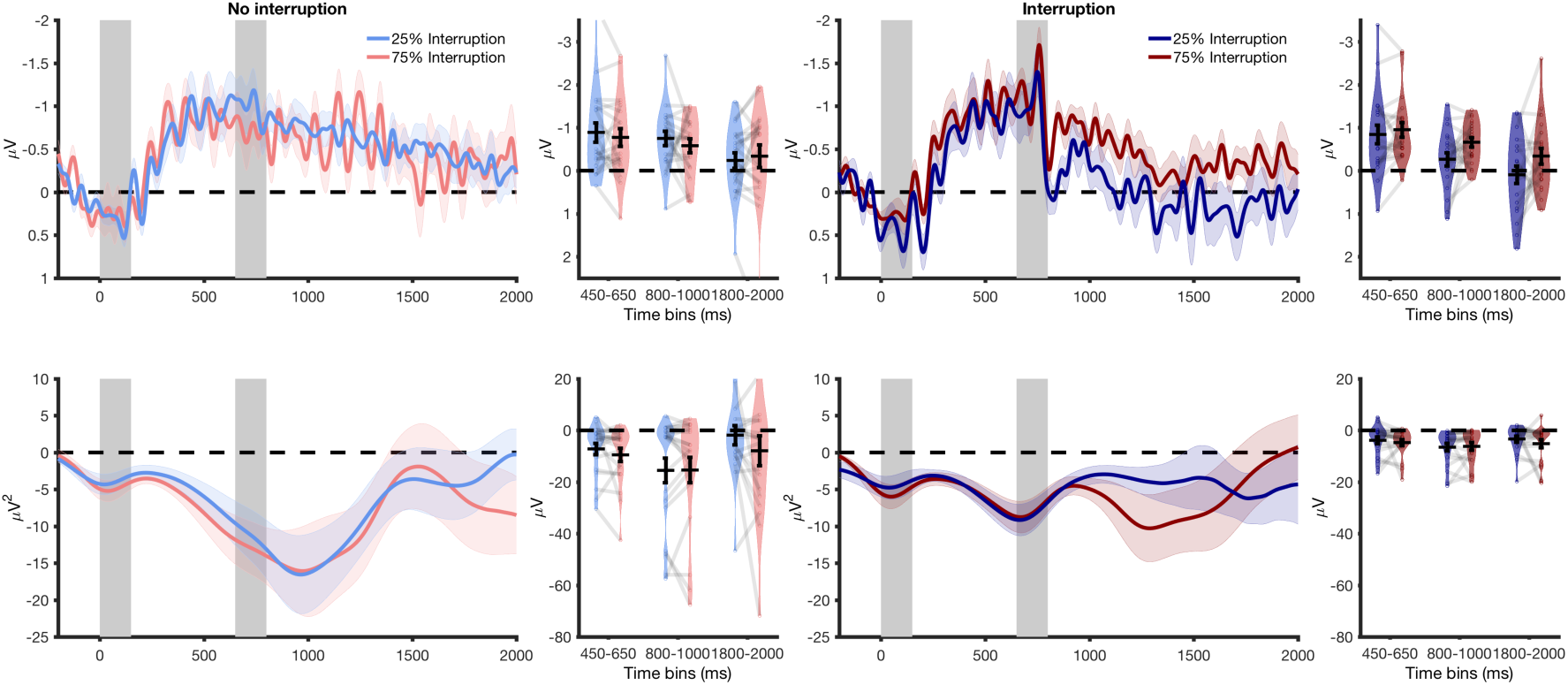
Contralateral delay activity and lateralized alpha power for Experiment 2. (Top) CDA amplitude over time. The first gray bar (timepoint 0-150) in both graphs represents when the memory array was on the screen. The second gray bar (timepoint 650-800) represents when the interrupters were on the screen in the Interruption condition. Light colored envelopes around each line represent standard error. (Bottom) Lateralized alpha power over time.

#### Post-interruption I (800-1,000 ms)

Immediately following the offset of interruption (800-1,000 ms), alpha power was significantly lateralized in all conditions, all one-sample t-tests p≤0.006. There was a significant main effect of Interruption on the strength of alpha lateralization: F(1,19)=6.530, p=0.019, η_p_^2^ = 0.256. For both High and Low probability trials, lateralized alpha power was stronger on trials without interruption (Low Probability M= -15.498, High Probability M= -15.368) than on trials with interruption (Low Probability M= -4.549, High Probability M= -4.966). No other effects were significant, p≥0.778.

#### Post-interruption II (1,800-2,000 ms)

By the end of the trial (1,800-2,000 ms), the influence of interruption on lateralized alpha power depended on the probability of interruption, significant interaction of Interruption and Probability, F(1,19)=6.365, p=0.021, η_p_^2^ = 0.251. The difference between lateralized alpha power in high and low probability trials is different with and without interruption. However, within High probability trials, follow-up t-tests revealed that there was not an alpha power suppression difference between trials that were interrupted and those that were not for either High (t(19)=-1.923, p=0.070) or Low (t(19)=1.077, p=0.295) probability trials. Also, alpha power suppression was not significantly different between High and Low probability trials for trials that were interrupted (t(19)=-1.278, p=0.217) or for those that were not interrupted (t(19)=1.362, p=0.189). The main effects of Interruption and Probability were also not significant, p≥0.472

### Conclusions

In Experiment 2, we found that behavioral performance was worse when participants were interrupted than when they were not interrupted, regardless of the probability of interruption. Once again, the neural results revealed that both the CDA and lateralized alpha power were negatively impacted by the interrupters, but these two signals had distinct time courses. Following interruption, the CDA sustained, but lateralized alpha power became less lateralized. In this time window, lateralization of alpha power did not depend on the probability of interruption. However, the amplitude of the CDA following interruption did depended on the probability of interruption – the CDA was larger when participants were expecting to be interrupted. By the end of the trial, the difference in lateralized alpha power between high and low probability trials reversed direction when comparing the difference between trials that were and were not interrupted. However, within high and within low probability trials, there was no difference between trials that were and were not interrupted. CDA by the end of the trial was no longer present, which may have been due to the extended retention interval. Probability had a clear effect on CDA amplitude, but the effect on lateralized alpha power was more ambiguous.

## General Discussion

Working memory maintains information so that it can be used despite momentary perturbations from task-irrelevant information. Here, we examined how memory representations that have already reached a stable state respond to visual interruption. As expected, we found a modest behavioral impact of interruption. Participants remembered significantly fewer items when they were interrupted than when they were not interrupted, but they performed above chance in all conditions. Despite a modest behavioral impact, task-irrelevant interruption produced substantial perturbations on two well-characterized EEG signals of working memory, lateralized alpha power and contralateral delay activity (CDA). Both lateralized alpha power, an index of sustained spatial attention, and CDA, an index of actively maintained working memory representations, were disrupted at certain points during the delay, but the time course of these perturbations varied. Lateralized alpha power results suggest that attention shifted towards baseline immediately following the interruption but had returned to the target positions by the end of the trial. By contrast, the CDA results suggest that working memory representations continued to persist following the interruption but was eliminated by the end of the trial. We additionally found that task expectancy modulated the timing and magnitude of these perturbations of working memory representations, suggesting that the brain’s response to task-irrelevant interruption is regulated by task context. The distinct time courses of and the influence of task context on lateralized alpha power and CDA have many interesting theoretical implications that future work can help elucidate.

### Neural response immediately following interruption

Sudden onsets of task-irrelevant interruption have been shown to capture attention when interrupters are visually salient (Bisley and Goldberg, 2003; Andrews et al., 2009; van Moorselaar et al., 2018). In our experiment, we used lateralized alpha power as an index of sustained spatial attention (Foster et al., 2016; Hakim et al., 2018). Following the onset of interruption, lateralized alpha power almost immediately shifted towards baseline. When lateralized alpha power is at baseline, it suggests that participants are no longer spatially attending the lateral memory items. Neural evidence from previous studies suggests that participants attend to the locations of interrupting stimuli (Bisley and Goldberg, 2003; van Moorselaar et al., 2018) because of attentional capture (Sawaki and Luck, 2012; Feldmann-Wüstefeld and Schubö, 2016). Thus, in the present study, participants presumably shifted their attention away from lateralized representations following the onset of task-irrelevant interruption to the centrally presented interrupters.

During this same time window, CDA remained robust and significantly above baseline. Previous research has shown that CDA is sensitive to trial-by-trial fluctuations in working memory performance and tracks the number of maintained object representations (Ikkai et al., 2010; Adam et al., 2015). Considering this, the robust CDA immediately following the onset of interruption suggests that object representations persist, at least momentarily, following the withdrawal of spatial attention to a new position. The presence of CDA and lack of lateralized alpha power immediately following interruption raises the long-standing theoretical question of whether object representations maintained in working memory can persist without sustained spatial attention. Previous research has suggested that spatial attention is a rehearsal mechanism that facilitates the maintenance of object representations held in working memory (Williams and Woodman, 2012). Additionally, the positions of object representations are maintained in working memory even when spatial information is completely irrelevant (Foster et al., 2017a). Together, these previous results suggest that spatial attention aids the maintenance of working memory information, but do not address whether working memory representations necessitate sustained spatial attention. In the present study, the robust CDA and lack of lateralized alpha power following the onset of interruption suggest that object representations maintained in working memory can persist without sustained spatial attention. Therefore, our results suggest that working memory representations may not necessitate sustained spatial attention. Nevertheless, working memory representations may still be volatile without sustained spatial attention, given that CDA goes to baseline by the end of trials with interruption.

### Neural activity at the end of interrupted trials

By the end of interrupted trials, CDA was no longer reliable, but alpha power became re-lateralized. This pattern of activity suggests that participants re-oriented their attention to the locations of the memoranda, but no longer maintained active working memory representations. If participants no longer maintain object representations by the end of the trial, how were they able to still perform the change detection task on interrupted trials (albeit worse than non-interrupted trials)? There are a few possible explanations.

One possible explanation for the absence of the CDA at the end of the trial but above chance behavioral performance is that performance on interrupted trials could rely on offline memory representations. Previous research has shown that information in working memory can be simultaneously maintained in both active and passive memory states (Mallett & Lewis-Peacock, 2018). Therefore, when actively maintained memory traces are no longer present, information could still be retrieved from an offline state. Research that has investigated retrieval of information from offline memory states has found that alpha power tracks information retrieved from long-term memory (Fukuda et al., 2016). Additionally, other research has suggested that attention can aid recall of information that would be otherwise unavailable to working memory (Murray et al, 2013). These findings dovetail with our results – at the end of interrupted trials, when information about the memoranda is required to respond to the probe, lateralized alpha power could be re-instated to aid the retrieval of information from offline memory storage, thereby bolstering behavioral performance.

Alternatively, the relatively good behavioral performance without CDA could be explained by other neural traces of actively maintained working memory representations that we are not measuring. The CDA is a coarse neural measure that compares activity contralateral and ipsilateral to memory items. Thus, it is not an exhaustive measure of working memory. More spatially global neural signals or more distributed patterns of activity, for example, could sustain following task-irrelevant interruption, and these signals could plausibly bolster behavioral performance. Regardless of the mechanism that preserves information about the memoranda, our results strongly suggest that actively maintained information is dynamically perturbed following task-irrelevant interruption.

### Modulation of CDA and alpha power by task demands

In Experiment 2, we varied task demands by interrupting participants on 75% (high) or 25% (low) of trials. Following interruption onset, we found the same pattern of result as Experiment 1; Lateralized alpha power shifted towards baseline while CDA persisted. However, the amplitude of the CDA varied as a function of task demands. When task demands were high, CDA amplitude was higher than when task demands were low. This suggests that participants were able to better protect working memory representations when they were expecting to be interrupted, and that task context is involved in how the brain responds to task-irrelevant interruption. On the other hand, the influence of task demands on lateralization of alpha power was more ambiguous. Our results suggest that spatial attention may be uniformly captured by interrupters initially regardless of expectation. However, during certain points in the trial, lateralization of alpha power may vary as a function of task demands. Therefore, the neural responses to interruption that we observed were affected both by both interruption and task demands. These results go hand-in-hand with previous research that has shown that distraction by salient irrelevant stimuli can be modulated by top-down control. For example, when a color singleton is presented on 20% of the trials, it slows down response times in a visual search task more than when it is presented on 50% of trials (Horstmann, 2005; Müller et al., 2009; Marini et al., 2013; Folk and Remington, 2015) because attention requires more time to be deployed to the relevant information when rare distractors appear (Töllner et al., 2012).

### Conclusions

In this set of experiments, we investigated the impact of task-irrelevant interruption on two dissociable neural signals, CDA, a neural index of actively maintained representations, and lateralized alpha power, an index of sustained spatial attention. By tracking these neural markers of working memory we were able to observe changes in active representations that would not be apparent from behavioral measures alone. Both CDA and lateralized alpha power were impacted by task-irrelevant information, yet had distinct time courses. Our results suggest that following interruption, lateralized visual representations of memoranda can stay active in working memory for a short period of time before they are lost. These representations do not recover by the end of the trial, and are presumed to be stored offline. By contrast, attention is directed away from the spatial location of memoranda immediately after the onset of the interruption but can recover later and may even contribute to the retrieval of information from offline storage. Thus, our results show that task-irrelevant interruption could motivate the transfer of information from active to passive storage. Moreover, the dissociation between CDA and lateralized alpha power further emphasizes that these neural markers distinctly contribute to the maintenance of information in working memory and may distinctly protect actively maintained memories from interruption.

## Acknowledgements

Research was supported by NIMH grant 5ROI MH087214-08 and Office of Naval Research grant N00014-12-1-0972.

